# Artificial Dendritic Neurons Enable Self-Supervised Temporal Feature Extraction

**DOI:** 10.1101/517888

**Authors:** Toshitake Asabuki, Tomoki Fukai

## Abstract

The brain identifies potentially salient features within continuous information streams to appropriately process external and internal temporal events. This requires the compression or abstraction of information streams, for which no effective information principles are known. Here, we propose conditional entropy minimization learning as the fundamental principle of such temporal processing. We show that this learning rule resembles Hebbian learning with backpropagating action potentials in dendritic neuron models. Moreover, networks of the dendritic neurons can perform a surprisingly wide variety of complex unsupervised learning tasks. Our model not only accounts for the mechanisms of chunking of temporal inputs in the human brain but also accomplishes blind source separation of correlated mixed signals, which cannot be solved by conventional machine learning methods, such as independent-component analysis.

**One Sentence Summary:** Neurons use soma-dendrite interactions to self-supervise the learning of characteristic features of various temporal inputs.

---

Cognitive functions of the brain entail modeling of externally or internally driven dynamic processes. For this modeling, the characteristic features of information streams must be identified by the brain at each stage of the hierarchical computation. Chunking or bracketing in such analyses underlies sensory, motor, and memory processing (1–5). However, the method by which neural circuits in the brain autonomously learn temporal features remains largely unclear. Here, we show that entropy minimization conditioned on a neuron’s own responses enables this learning in an unsupervised fashion. We demonstrate that networks of artificial dendritic neurons can self-supervise the learning of spatiotemporal firing patterns that are repeatedly evoked in upstream neurons. This model enables the learning of a surprisingly wide variety of tasks, including the chunking of temporal inputs, the formation of orientation maps, and even the blind source separation (BSS) of correlated mixed signals. Although BSS has been extensively studied for independent signals (6–8), no effective methods except for semi-supervised methods are known for the processing of correlated signals (9).

Our model entails the learning of temporal features of an input based on a novel learning rule, which we term “self-conditioned entropy minimization (SCEM).” In short, SCEM categorizes temporal inputs by minimizing variations in neuronal responses to a given set of external inputs. The variation will be minimal when a neuron responds similarly to similar inputs. To achieve this, SCEM learns to self-generate appropriate teaching signals. Figure 1A shows a biology-inspired implementation of SCEM in a two-compartment spiking neuron model (see Supplementary Materials for mathematical details). In short, activity in the dendritic compartment, driven by external inputs, predicts somatic spike responses. This division of labor between somatic and dendritic compartments has been explored in a neuron model for supervised learning, with a teaching signal given to the soma (10). Unlike the previous model, our neuron model performs unsupervised learning by feeding the somatic response back to the dendrite to train dendritic synapses. Although the underlying biological mechanisms require further clarification, backpropagating action potentials may provide the feedback signal for this self-supervision in cortical pyramidal neurons (11, 12). Our learning rule (Eq. 19 in Supplementary Materials) looks similar to the maximum likelihood estimation (13), a well-studied framework of supervised learning. However, there is a conceptual difference between them. In the maximum likelihood estimation, the target data distribution (somatic activity) is provided externally as teaching signals. By contrast, our model learns the simultaneous distributions of input and output data without teaching signals. The consistency between the two data sets constrains the self-supervised learning, thereby avoiding a redundant or an overly simplistic categorization of temporal inputs. Although SCEM fits particularly well with dendritic neurons, the principle is generic and applicable to a broad range of information processing systems.

**Fig. 1.**
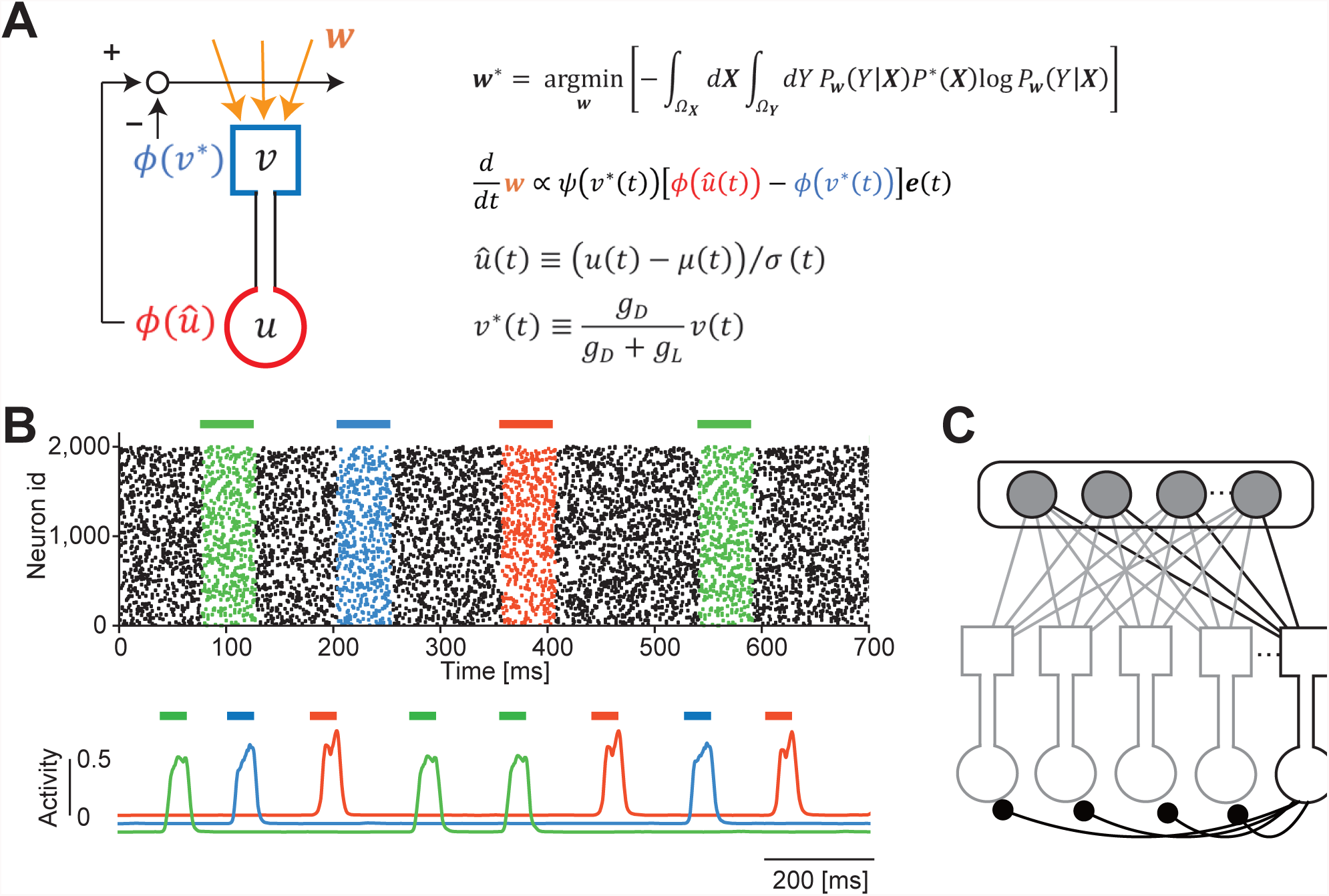
Unsupervised learning in two-compartment neurons. (A) The model neuron consists of somatic and dendritic compartments and undergoes SCEM learning. The dendritic component receives Poisson spike trains, and the somatic membrane potential is given as an attenuated version of the dendritic membrane potential. Output of the soma backpropagates to dendritic synapses as a self-teaching signal. Learning stops when the dendrite minimizes the error between its prediction and the actual somatic firing rate. (B) Three frozen spatiotemporal patterns (red, blue, and green) were repeated as irregular spike trains from 2,000 input neurons (top). Three dendritic neurons selectively responded to one of the repeated patterns after learning (bottom). (C) A competitive network used in all of the present tasks. The input layer consists of Poisson spiking neurons, and the output layer comprises the dendritic neuron models. Ten output neurons were connected with all-to-all inhibitory synapses modifiable by iSTDP.

As shown in Fig. 1B (top), presynaptic spike trains intermittently repeated three fixed spatiotemporal patterns with equal probabilities of occurrence. The learning of repeated temporal input patterns is crucial for various cognitive functions such as language acquisition (14, 15) and motor sequence learning (1–3, 16). A single neuron learned to respond selectively to one of the input patterns (Fig. 1B, bottom), with approximately equal probabilities for the patterns among the trials, although it responded to more than one input pattern in some cases (Fig. S1). Cortical neurons actually have the ability to discriminate simple temporal inputs (17). Next, we considered a competitive network of two-compartment model neurons receiving similar presynaptic spike trains (Fig. 1C). Recurrent inhibitory connections among these neurons were modifiable by inhibitory spike timing-dependent plasticity (iSTDP; Fig. S2A). The postsynaptic neurons self-organized into three neuron ensembles, each detecting one of the input activity patterns (Fig. S2B). iSTDP enabled mutual inhibition between the neural ensembles (Fig. S2C). The strength of lateral inhibition required adjustment, as inhibition that was too strong (Fig. S3A, B) or too weak (Fig. S3C, D) eliminated chunk-specific cell assemblies. These results may explain how humans can detect frozen noise patterns repeated within noisy auditory signals. The regularization parameter *γ* (see Materials and Methods) must also be in an appropriate range to enable the unsupervised learning of chunk-specific cell assemblies, as values that are too large suppress all neural responses and those that are too small do not generate selective responses to chunks (Fig. S4).

The ability of the network model to learn was assessed with various types of biological noise. Background presynaptic spikes degraded the performance as the signal-to-noise ratio decreased (Fig. S5A), whereas learning was optimal at finite noise levels with synaptic transmission failure (Fig. S5B) and with jitters in presynaptic spike timing (Fig. S5C). We speculate that this disparity may reflect the different noise structures. Background spikes were not correlated with the repeated input patterns and merely contaminated the signals, whereas the noise patterns from transmission failures and timing jitters yielded noise that was correlated and thus enhanced the sampling for learning. Although presynaptic noise may induce a regularization effect during learning (18), this likely did not occur in our model network, as not all types of presynaptic noise improved the learning.

The network model is capable of learning repeated patterns in various information streams. To show this, we applied random sequences of three chunks comprising four characters each (Fig. 2A) to a network model with 10 output neurons and 1,000 input neurons. Each input neuron generated a 30 ms 10 Hz burst in response to a randomly assigned preferred character (Fig. 2B). This resulted in the formation of three neuron ensembles that selectively responded to the chunks (Fig. 2C). We conducted a principal-component analysis to study the low-dimensional dynamics of output neurons, which revealed the emergence of the three chunks after learning (Fig. 3D). However, the word segmentation shown above is not difficult for other methods as well (19). Therefore, we next tested the same model with more complex input sequences generated by a random walk on a graph with a community structure, where the connection of each node to the other four occurred with an equal probability of 0.25 (Fig. 2E). The detection of this community structure is easy for human subjects but difficult by construction for the conventional machine learning methods that rely on surprise signals, such as those with nonuniform transition probabilities (4). To our surprise, each output neuron easily learned to respond selectively to members within its community (Fig. 2F).

**Fig. 2.**
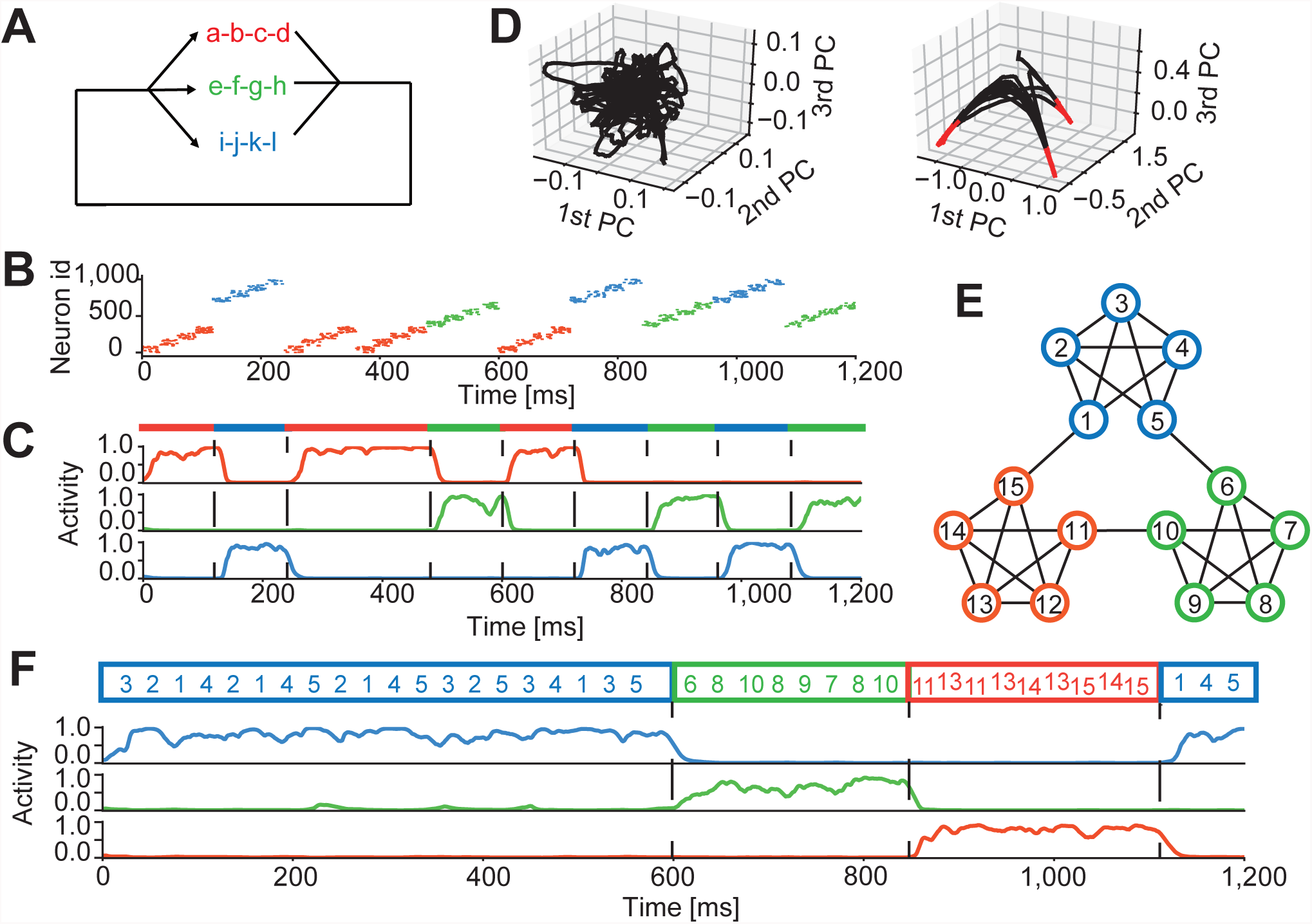
Segmentation and concatenation of various sequences. (A) Three chunks (a-b-c-d [red], e-f-g-h [green], and i-j-k-l [blue]) repeatedly appeared in the input sequence with equal probabilities. (B) Each input neuron fired at 10 Hz to encode one of the chunks. Neurons were sorted according to their preferred stimuli. (C) Typical normalized responses of three output neurons are shown after learning. Colors indicate the epochs of the corresponding chunks. (D) Responses of output neurons were projected onto the three leading principal-component (PC) vectors before (left) and after (right) learning. Epochs of high normalized responses (*f* > 0.8 in all neurons) are indicated in red. (E) The input sequence represented a random walk with uniform transition probabilities on a graph with community structure (modified from reference 4). (F) Normalized responses of three output neurons to input sequences defined in panel E are shown.

**Fig. 3.**
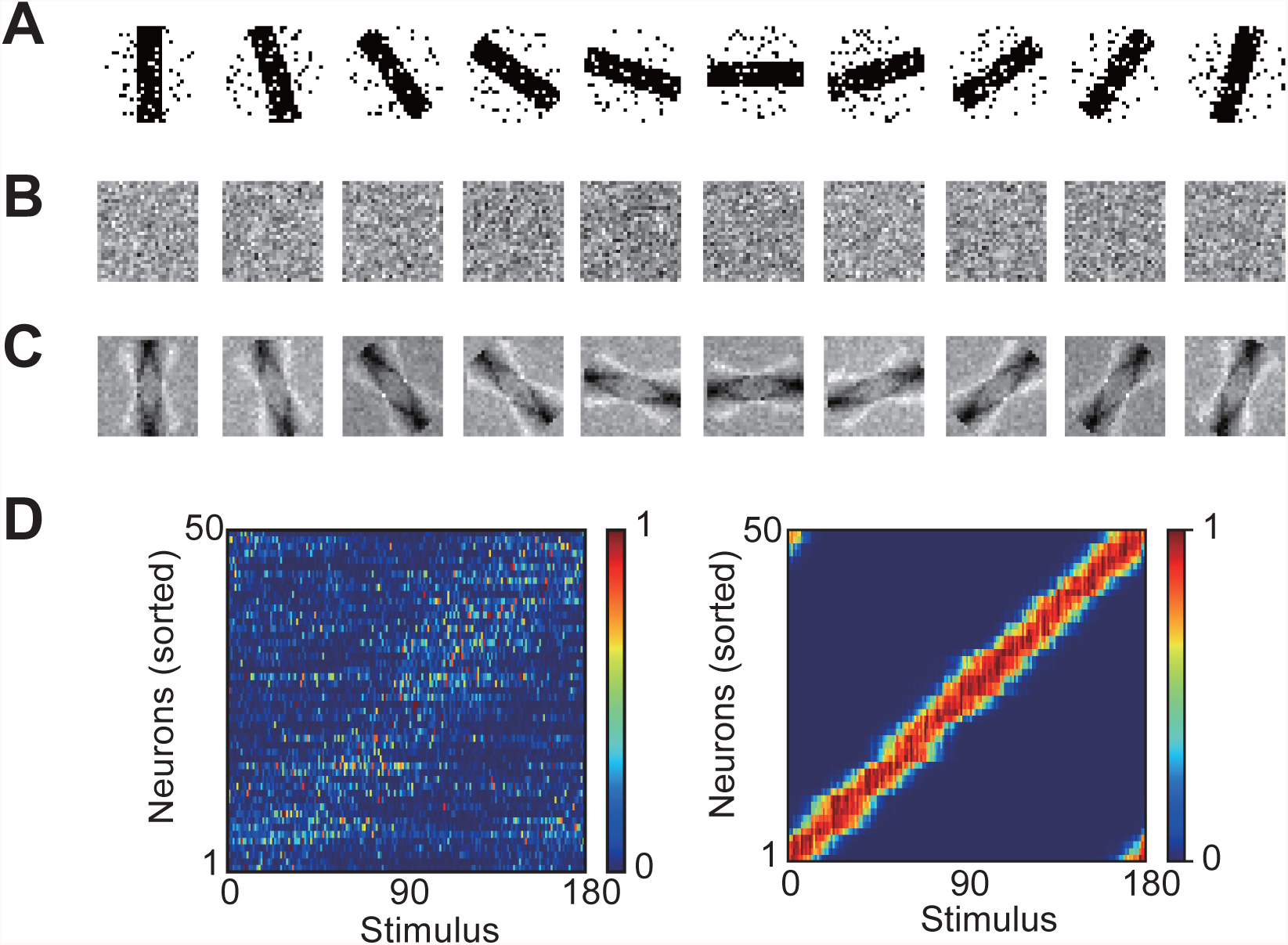
Learning an orientation tuning map. (A) Examples of noisy images of oriented bars used for the training. Each image was presented for 40 ms in a random order with intervals of 30 ms between images. (B and C) The feedforward synaptic weights before and after learning are shown for the example stimuli shown in panel A. (D) The responses of all dendritic neurons before (left) and after (right) learning are shown. The neurons were sorted according to the onset times of responses to their preferred stimuli. See Supplementary Text for further details on the simulations.

The network model also learns the static features of input when they are repeatedly shown in a temporal sequence. A random sequence of noisy images of oriented bars presented for 40 ms at 30 ms intervals was applied to the model (Fig. 3A). The output neurons, which initially had no preferred orientation (Fig. 3B), developed well-defined preferences for specific orientations after learning (Fig. 3C), resembling a visual orientation map (Fig. 3D) (20, 21). Because all sensory features, either static or dynamic, arrive at the brain in sequence, temporal processing is potentially important for the formation of feature detection maps from continuous sensory streams.

These results demonstrate that the SCEM successfully chunks a variety of temporal inputs by automatically identifying repeated temporal input patterns. The question then arises whether this ability of the SCEM enables learning of other types of sequence processing tasks. Sequence processing also involves the blind separation of signals within mixtures from multiple sources. BSS is an extensively studied problem in auditory processing (6–8), but the various methods that have been proposed are effective only if individual signals are independent. To our knowledge, there are no effective methods for separating mixtures of dependent or correlated signals. We applied SCEM to sound mixtures from two music instruments (Audio S1), i.e., a bassoon and a clarinet (Bach10 Dataset (22); Audios S2 and S3), playing their respective parts of the same score (Fig. 4A); thus the two sound sources are correlated. These mixtures of signals were encoded as irregular spike trains (Fig. 4B), which in turn were applied to output neurons. After training, these neurons self-organized into two subgroups, each responding to one of the true sources (Fig. 4C). The original sounds were then decoded from the average firing rates of these subgroups (Audios S4 and S5), though some high-frequency components were lost due to the low-pass filtering effect corresponding to membrane dynamics (Fig. S6). By contrast, BSS was poor via an independent-component analysis (Audio S6).

**Fig. 4.**
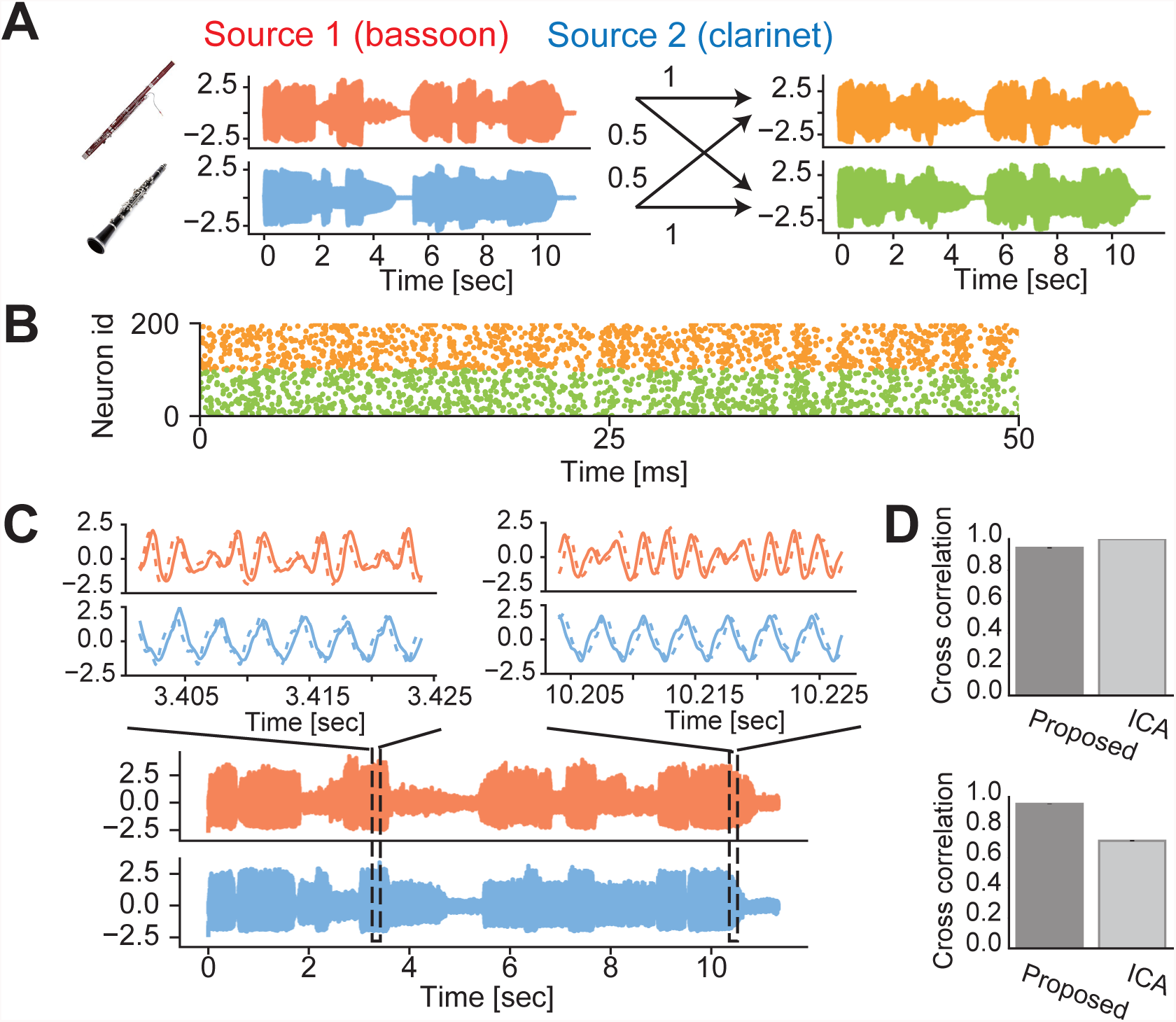
BSS of dependent auditory streams. (A) Sound waveforms of a bassoon and a clarinet (left) were linearly transformed to two mixture signals (right). The diagonal and off-diagonal elements of the mixing matrix were 1 and 0.5, respectively. (B) Nonstationary Poisson spike trains of 200 input neurons (out of the total 500) are shown. The instantaneous firing rates were proportional to the amplitudes of the mixed signals normalized between 0 Hz and 10 Hz. Each input neuron encodes either of the two mixed signals. (C) Separated waveforms (bottom) are shown together with magnified versions (top, solid) and true sources (top, dashed). The waveforms were averaged over 20 trials with different realizations of input spike trains and the same initial weights. (D) Cross-correlations between the separated and true sources are compared between our model and independent-component analysis (ICA) for independent (top) and dependent (bottom) auditory signals (see Supplementary Materials). Error bars show SDs (invisible).

Mutual information maximization (MIM) has often been hypothesized to describe the transfer of information between neurons (23), and Hebbian synaptic plasticity may approximately follow MIM (24). However, the MIM principle ultimately implies that messages are faithfully copied at all layers of hierarchical processing. Furthermore, MIM does not account for the compression or abstraction of sensory input to the brain.

Our learning rule minimizes the entropy associated with the conditional probability of neuronal output for a given input. The rule enabled mutually inhibiting dendritic neurons to learn the repetition of spatiotemporal activity patterns on a slow timescale (typically, several tens to several hundreds of milliseconds). While the aim of many previous methods for chunking is to predict the input sequence (25, 26), our model entails a novel principle in which a neural system learns to predict its own responses to input. To this end, the SCEM minimizes the conditional entropy of output data to produce a predictable low-dimensional representation of high-dimensional input data. This learning continues until there is agreement between the somatic output and dendritic input regarding the low-dimensional features (i.e., chunks). We previously used paired reservoir computing for chunking (but not for BSS), in which two recurrent networks supervise each other to mimic the partner’s responses to a common temporal input (27). The present model outperforms the previous one, but the two models share a fundamental computational principle, namely, self-consistency between input and output data. The SCEM, on the other hand, differs from autoencoders that compress input information in hidden layers. The compression rate is much higher in our model than in autoencoders, because input sequences cannot be faithfully reconstructed from chunked pieces. Despite resembling methods for learning the probabilistic structure of input data (10, 13) and the fact that the information bottleneck compresses data while maintaining mutual information to some degree (28), the SCEM differs from these and other methods aimed at learning the likelihood of the input data distribution.

In sum, our model not only performs chunking but also achieves BSS from mixtures of correlated signals. It is surprising that simple neural networks with identical circuit structures can perform these seemingly different tasks. Such a multifunctional model was previously unknown in learning information streams.

## Supporting information

AudioS1

AudioS2

AudioS3

AudioS4

AudioS5

AudioS6

## Acknowledgments

The authors express their sincere thanks to Shun-ichi Amari for stimulating discussion about the learning rule and to Shigeyoshi Fujisawa and Joshua Johansen for their valuable comments on our manuscript.

## Author contributions

T.F. and T.A. designed the study and wrote the manuscript, and T.A. performed numerical simulations.

## Data and materials availability

The computer code used in this study is available from the authors upon request.

## Supplementary Materials

Materials and Methods

Supplementary Text

Figures S1–S6

Audio Files S1–S6

## References

1. N. Fujii, A. M. Graybiel, Representation of action sequence boundaries by macaque prefrontal cortical neurons. Science. 301, 1246–1249 (2003).

2. X. Jin, R. M. Costa, Start/stop signals emerge in nigrostriatal circuits during sequence learning. Nature. 466, 457–462 (2010).

3. X. Jin, F. Tecuapetla, R. M. Costa, Basal ganglia subcircuits distinctively encode the parsing and concatenation of action sequences. Nat Neurosci. 17, 423–430 (2014).

4. A. C. Schapiro, T. T. Rogers, N. I. Cordova, N. B. Turk-Browne, M. M. Botvinick, Neural representations of events arise from temporal community structure. Nat Neurosci. 16, 486–492 (2013).

5. J. M. Zacks, T. S. Braver, M. A. Sheridan, D. I. Donaldson, A. Z. Snyder, J. M. Ollinger, R. L. Buckner, M. E. Raichle, Human brain activity time-locked to perceptual event boundaries. Nat Neurosci. 18, 449–455 (2001).

6. P. Comon, Independent component analysis, A new concept? Signal Processing. 36, 287–314 (1994).

7. S. Amari, J. F. Cardoso, Blind source separation – semiparametric statistical approach. IEEE Transactions on Signal Processing. 45, 2692–2700 (1997).

8. A. Hyvärinen, E. Oja, Independent component analysis: Algorithms and applications. Neural Networks. 13, 411–430 (2000).

9. H. Kameoka, L. Li, S. Inoue, S. Makino, https://arxiv.org/abs/1808.00892 (2018).

10. R. Urbanczik, W. Senn, Learning by the dendritic prediction of somatic spiking. Neuron. 81, 521–528 (2014).

11. M. Larkum, A cellular mechanism for cortical associations: An organizing principle for the cerebral cortex. Trends Neurosci. 36, 141–151 (2013).

12. M. E. Larkum, J. J. Zhu, B. Sakmann, A new cellular mechanism for coupling inputs arriving at different cortical layers. Nature. 398, 338–341 (1999).

13. J. P. Pfister, T. Toyoizumi, D. Barber, W. Gerstner, Optimal Spike-Timing Dependent Plasticity for Precise Action Potential Firing. Neural Computation. 18, 1318–1348 (2006).

14. M. Buiatti, M. Peña, G. Dehaene-Lambertz, Investigating the neural correlates of continuous speech computation with frequency-tagged neuroelectric responses. Neuroimage. 44, 509–519 (2009).

15. T. Q. Gentner, K. M. Fenn, D. Margoliash, H. C. Nusbaum, Recursive syntactic pattern learning by songbirds. Nature. 440, 1204–1207 (2006).

16. K. S. Smith, A. M. Graybiel, A dual operator view of habitual behavior reflecting cortical and striatal dynamics. Neuron. 79, 361–374 (2013).

17. T. Branco, B. A. Clark, M. Häusser, Dendritic discrimination of temporal input sequences in cortical neurons. Science. 329, 1671–1675 (2010).

18. C. M. Bishop, Training with noise is equivalent to Tikhonov regularization. Neural Comput. 7, 108–116 (1995).

19. P. Pierre, What Mechanisms Underlie Implicit Statistical Learning? Transitional Probabilities Versus Chunks in Language Learning. Topics in Cognitive Science. DOI: 10.1111/tops.12403 (2018).

20. D. H. Hubel, T. N. Wiesel, Receptive fields of single neurons in the cat’s striate cortex. J Physiol. 148, 574–591 (1959).

21. B. A. Olshausen, D. J. Field, Emergence of simple-cell receptive field properties by learning a sparse code for natural images. Nature. 381, 607–609 (1996).

22. Z. Duan, B. Pardo, Soundprism: an online system for score-informed source separation of music audio. IEEE Journal of Selected Topics in Signal Process. 5, 1205–1215 (2011).

23. F. Rieke, D. Warland, R. de Ruyter van Stevenick, W. Bialek, Spikes (Cambridge, MA: Massachusetts Institute of Technology, 1997).

24. R. Linsker, Perceptual neural organization: some approaches based on network models and information theory. Annu Rev Neurosci. 13, 257–281 (1990).

25. C. Wacongne, J. P. Changeux, S. Dehaene, A Neuronal Model of Predictive coding accounting for the mismatch negativity. Journal of Neurosci. 32, 3665–3678 (2012).

26. S. J. Kiebel, K. V. Kriegstein, D. Daunizeau, L. J. Friston, Recognizing sequences of sequences. PLoS Comput Biol. 6, e1000464 (2009).

27. T. Asabuki, N. Hiratani, T. Fukai, Interactive reservoir computing for chunking information streams. PLoS Comput Biol. 14, e1006400 (2018).

28. N. Tishby, F. C. Pereira, W. Bialek, Proc. of the 37th annual Allerton Conference on Communication, Control, and Computing, 368–377 (1999).

29. L. Wiskott, T. J. Sejnowski, Slow feature analysis: unsupervised learning of invariances. Neural Comput, 14, 715–770 (2002).

30. A. P. Dempster, N. M. Laird, D. B. Rubin, Maximum likelihood from incomplete data via the EM algorithm. Journal of the Royal Statistical Society, Series B. 39, 1–38 (1977).

31. A. Hyvärinen, Fast and robust fixed-point algorithms for independent component analysis. IEEE Transactions on Neural Networks. 10, 626–634 (1999).

32. J. T. Lu, C. Y. Li, J. P. Zhao, M. M. Poo, X. H. Zhang, Spike-timing-dependent plasticity of neocortical excitatory synapses on inhibitory interneurons depends on target cell type. J Neurosci 27, 9711–9720 (2007).

