## Supplementary Materials for "Artificial Dendritic Neurons Enable Self-Supervised Temporal Feature Extraction"

**This PDF file includes:**

Methods
Supplementary Text
Figs. S1 to S6
Captions for Audios S1 to S6

**Other Supplementary Materials for this manuscript include the following:**

Audios S1 to S6

### Methods

Neural network model

Each output neuron has two compartments, i.e., somatic and dendritic compartments. The dendritic membrane potential of output neuron $i \in \{1,2,\dots,N_{\text{out}}\}$ is calculated as

$$v_i(t) = \sum_j w_{ij} e_j(t), \tag{1}$$

where $w_{ij}$ is the weight of synapse between output neuron $i$ and input neuron $j$. The somatic activity integrates the dendritic potential, and it evolves as

$$\dot{u}_i(t) = -\frac{1}{\tau} u_i(t) + g_D[-u_i(t) + v_i(t)] - \sum_j G_{ij} f_j(t), \tag{2}$$

where $\tau = 15$ ms and the conductance between the two compartments $g_D = 0.7$. The last term describes lateral inhibition with synaptic weights $G_{ij}$ ($\geq 0$). We assume that the soma of neuron $i$ generates a Poisson spike train with the instantaneous firing rate

$$f_i(t) = \phi(u_i(t)), \tag{3}$$

in terms of a nonlinear response function

$$\phi(x) = \phi_0\left[1 + \exp\left(\beta(-x + \theta)\right)\right]^{-1}, \tag{4}$$

with $\beta = 5$. Values of the parameters $\phi_0$ and $\theta$ are task-depend and described later.

Input neuron $i \in \{1,2,\dots,N_{\text{in}}\}$ generates a Poisson spike train

$$X_i(t) = \sum_q \delta(t - t_{i,q}), \tag{5}$$

where $\delta$ is the Dirac' delta function and $t_{i,q}$ denotes the time of the $q$-th spike of input neuron $i$. The presynaptic spikes induce the following synaptic current $I_i(t)$:

$$\tau_{\text{syn}} \dot{I}_i = -I_i + \frac{1}{\tau} X_i\,, \tag{6}$$

where the synaptic time constant $\tau_{\text{syn}} = 5$ ms. The synaptic currents in turn evoke postsynaptic potential (PSP) $e_i(t)$ as

$$\dot{e}_i = -\frac{e_i}{\tau} + I_i. \tag{7}$$

Self-constrained entropy minimization (SCEM)

To extract the characteristic features of temporal signal, our model compresses the high dimensional data of input sequence onto a low dimensional manifold of neural dynamics. Our model performs this by modifying the weights of dendritic synapses to minimize the conditional entropy of the entire network activity. The conditional entropy is defined as

$$\mathcal{H} = -\langle \log P_{\boldsymbol{w}}(\boldsymbol{Y}|\boldsymbol{X}) \rangle_{P_{\boldsymbol{w}}(\boldsymbol{Y}|\boldsymbol{X})P^*(\boldsymbol{X})}, \tag{8}$$

where $P^*(\boldsymbol{X})$ is the true distribution of input spike trains and $\boldsymbol{Y} = [Y_1, \cdots, Y_{N_{\text{out}}}]$ represents output spike trains during the trial $T$, with $Y_i$ being the spike train of output neuron $i$. The bracket represents the averaging $\langle q(\boldsymbol{X}, \boldsymbol{Y}) \rangle_{P_{\boldsymbol{w}}(\boldsymbol{Y}|\boldsymbol{X})P^*(\boldsymbol{X})} = \int_{\Omega_X} d\boldsymbol{X} \int_{\Omega_Y} d\boldsymbol{Y}\, q(\boldsymbol{X}, \boldsymbol{Y})\, P_{\boldsymbol{w}}(\boldsymbol{Y}|\boldsymbol{X})P^*(\boldsymbol{X})$, where the integration should be taken over all possible combinations of input and output spike trains ($\Omega_{\boldsymbol{X}}$ and $\Omega_{\boldsymbol{Y}}$, respectively). Because the entropy is conditioned on the probability distribution of the system's own outputs, we may term $\mathcal{H}$ "the self-conditioned entropy". A similar objective function was considered in a previous modeling study, but the entropy was conditioned on the probability distribution of external teaching signals (*10*).

If the individual output neurons fire independently (this only holds approximately in the presence of lateral inhibition), $P(\boldsymbol{Y}|\boldsymbol{X}) = \prod_i P_{\boldsymbol{w}_i}(Y_i|\boldsymbol{X})$, where $P_{\boldsymbol{w}_i}(Y_i|\boldsymbol{X})$ is the probability density of output spike train $Y_i$ for given weight vector $\boldsymbol{w}_i = [w_{i1}, \cdots w_{iN_{\text{in}}}]$ and input spike train $\boldsymbol{X}$. Then, $\langle \cdot \rangle_{P_{\boldsymbol{w}}(\boldsymbol{Y}|\boldsymbol{X})P^*(\boldsymbol{X})} = \int_{\Omega_X} d\boldsymbol{X} \int_{\Omega_Y} \cdots \int_{\Omega_Y} \prod_i dY_i \cdot P_{\boldsymbol{w}_i}(Y_i|\boldsymbol{X})P^*(\boldsymbol{X})$, and we can derive the following upper bound for the self-conditioned conditioned entropy:

$$\begin{aligned}
\mathcal{H} &\leq -\int_{\Omega_X} d\boldsymbol{X} \int_{\Omega_Y} \cdots \int_{\Omega_Y} \prod_i dY_i \log\left[\prod_j P_{\boldsymbol{w}_j}(Y_j|\boldsymbol{X})\right] \prod_k P_{\boldsymbol{w}_k}(Y_k|\boldsymbol{X})\ P^*(\boldsymbol{X}) \\
&= -\sum_j \int_{\Omega_X} d\boldsymbol{X} \left[\prod_{i \neq j} \int_{\Omega_Y} dY_i\, P_{\boldsymbol{w}_i}(Y_i|\boldsymbol{X})\right] \int_{\Omega_Y} dY_j\ P^*(\boldsymbol{X}) P_{\boldsymbol{w}_j}(Y_j|\boldsymbol{X}) \log P_j(Y_j|\boldsymbol{X}) \\
&= -\sum_j \int_{\Omega_X} \int_{\Omega_Y} P_{\boldsymbol{w}_j}(\boldsymbol{X}, Y_j) \log P_{\boldsymbol{w}_j}(Y_j|\boldsymbol{X}) d\boldsymbol{X} dY_j.
\end{aligned} \tag{9}$$

Here, we have used the relationship $P_{\boldsymbol{w}_i}(\boldsymbol{X}, Y_i) = P_{\boldsymbol{w}_i}(Y_i|\boldsymbol{X})P^*(\boldsymbol{X})$. From Eq. 9, we obtain the optimal weight matrix $\boldsymbol{w}^*$ that minimizes the upper bound as

$$\boldsymbol{w}^* = \underset{\boldsymbol{w}}{\operatorname{argmax}} \sum_i \langle \log P_{\boldsymbol{w}_i}(Y_i|\boldsymbol{X}) \rangle_{P_{\boldsymbol{w}_i}(\boldsymbol{X}, Y_i)}. \tag{10}$$

Note that the $i$-dependence of $\boldsymbol{w}_i$ arises from activity-dependent modifications of recurrent inhibitory connections among output neurons (see equation 2). These inhibitory connections were modifiable by STDP (see Fig. S2C).

We next calculate the conditional probability, $P_{\boldsymbol{w}_i}(Y_i|\boldsymbol{X})$. We assumed that an attenuated version of dendritic potential $v_i^*(t)$ well describes the somatic membrane potential as $u(t) \approx v_i^*(t) = \alpha v_i(t)$ , where the degree of attenuation $\alpha = g_D/(g_D + g_L)$ (*10*). Noting that the probability of having a spike at $t' \in Y_i$ is $\prod_{t' \in Y_i} \phi\big(u_i(t')\big)$ and that of having no spikes at elsewhere is $\exp\left(-\int_0^T \phi\big(u_i(s)\big)ds\right)$, we can approximate $P_{\boldsymbol{w}_i}(Y_i|\boldsymbol{X})$ as

$$P_{\boldsymbol{w}_i}(Y_i|\boldsymbol{X}) \approx \left[\prod_{t' \in Y_i} \phi\big(v_i^*(t')\big)\right] \exp\left(-\int_0^T \phi\big(v_i^*(t)\big)dt\right)$$
$$= \exp\left(\int_0^T \left[\log\Big(\phi\big(v_i^*(t)\big)\Big) Y_i(t) - \phi\big(v_i^*(t)\big)\right] dt\right). \quad (11)$$

Constraints on somatic activity

The optimization problem described in the previous subsection has a trivial solution $\boldsymbol{w} = \boldsymbol{0}$. In this case, the firing rate of output neuron become always zero regardless of input $X$. To avoid the trivial solution, we adopted the empirical methodology that was proposed in the slow-feature analysis of temporal input (*29*). Namely, we renormalized the somatic membrane potential as follows:

$$\hat{u}_i(t) \equiv \big(u_i(t) - \mu_i(t)\big)/\sigma_i\,(t), \quad (12)$$

where $\mu_i(t)$ and $\sigma_i(t)$ are the mean and variance of the membrane potential

$$\mu_i(t) = \frac{1}{t_0}\int_{t-t_0}^{t} u_i(t')\,dt', \quad (13)$$

$$\sigma_i(t) = \sqrt{\frac{1}{t_0}\int_{t-t_0}^{t} u_i(t')^2\,dt' - \mu_i(t)^2}, \quad (14)$$

over a sufficiently long period $t_0$. Then, we replaced Eq. 3 with the following equation:

$$f_i(t) = \phi\big(\hat{u}_i(t)\big). \quad (15)$$

Using the renormalized somatic activity, we can avoid the saturation of output firing rate (as the mean of $\hat{u}_i(t)$ is constrained to be zero) and obtain fluctuations of order unity around the mean.

Optimal learning rule for SCEM

Now we derive the learning rule for synaptic weights. Because the conditional probability $P_{\boldsymbol{w}_i}(Y_i|\boldsymbol{X})$ is not explicitly known, the direct maximization of the expectation value $\sum_i \langle \log P_{\boldsymbol{w}_i}(Y_i|\boldsymbol{X}) \rangle_{P_{\boldsymbol{w}_i}(\boldsymbol{X},Y_i)}$ is difficult. Alternatively, we employed an EM-like optimization method (*29*). In the E-step, we sampled the somatic spike trains for given input from the joint distribution $\hat{P}_i(\boldsymbol{X}, Y_i) = \hat{P}_i(Y_i|\boldsymbol{X})P^*(\boldsymbol{X})$, where $\hat{P}_i(Y_i|\boldsymbol{X})$ is a constrained version of $P_{\boldsymbol{w}_i}(Y_i|\boldsymbol{X})$ and was obtained by sampling spike trains generated by the renormalized version of somatic activity.

In the M-step, we maximized the expectation value $\sum_i \langle \log P_{\boldsymbol{w}_i}(Y_i|\boldsymbol{X}) \rangle_{\hat{P}_i(\boldsymbol{X},Y_i)}$ over the fixed distribution $\hat{P}_i(\boldsymbol{X}, Y_i)$ obtained in the preceding E-step. Thus, weight changes were calculated by the stochastic gradient ascent as

$$\begin{aligned}\Delta w_{ij} &\propto \frac{\partial}{\partial w_{ij}} \sum_{i'} \left\langle \log P_{\boldsymbol{w}_{i'}}(Y_{i'}|\boldsymbol{X}) \right\rangle_{\hat{P}_{i'}(\boldsymbol{X},Y_{i'})} \\ &= \left\langle \int_0^T \frac{\partial}{\partial w_{ij}} \left[ \log\left(\phi\left(v_i^*(t)\right)\right) Y(t) - \phi\left(v_i^*(t)\right)\right] dt \right\rangle_{\hat{P}_i(\boldsymbol{X},Y_i)} \\ &= \left\langle \int_0^T \frac{\partial \log\left(\phi\left(v_i^*(t)\right)\right)}{\partial w_{ij}} \left[Y(t) - \phi\left(v_i^*(t)\right)\right] dt \right\rangle_{\hat{P}_i(\boldsymbol{X},Y_i)} . \end{aligned} \tag{16}$$

Note that the identity $d\phi(x)/dx = \phi(x) d\log\phi(x)/dx$ was used in deriving the last expression. Since $v_i^*(t) = \alpha \sum_j w_{ij} e_j(t)$, this yields the local synaptic learning rule

$$\Delta w_{ij} \propto \left\langle \int_0^T \psi\left(v_i^*(t)\right) \left[Y(t) - \phi\left(v_i^*(t)\right)\right] e_j(t) dt \right\rangle_{\hat{P}_i(\boldsymbol{X},Y_i)} , \tag{17}$$

where the function $\psi(x)$ is defined as

$$\psi(x) = \frac{d}{dx} \log\left(\phi(x)\right). \tag{18}$$

In all of the simulation, we replaced the output spike trains $Y(t)$ with the firing rate $\phi\left(\hat{u}_i(t)\right)$, and employed the online learning rule given as

$$\dot{w}_{ij}(t) = \eta \left\{ \psi\left(v_i^*(t)\right)\left[\phi\left(\hat{u}_i(t)\right) - \phi\left(v_i^*(t)\right)\right] e_j(t) - \gamma w_{ij} \right\}, \tag{19}$$

where $\eta$ is the learning rate. In the above equation, a regularization term $-\gamma w_{ij}$ was introduced to prevent a diverging growth of synaptic weights. The parameter $\gamma$ controls the strength of regularization and was adjusted in a task-dependent manner. The initial values of $\boldsymbol{w}$ were generated by a Gaussian distribution with mean zero and standard deviation $1/\sqrt{N_{\text{in}}}$.

Inhibitory plasticity

We modified lateral inhibitory connections through a symmetric anti-Hebbian spike-timing-dependent plasticity (STDP): If a pair of presynaptic and postsynaptic spikes occur at the times $t_{\text{pre}}$ and $t_{\text{post}}$, respectively, weight changes were calculated as

$$\Delta G_{ij} = C_p \exp\left(-\frac{|t_{\text{pre}} - t_{\text{post}}|}{\tau_p}\right) - C_d \exp\left(-\frac{|t_{\text{pre}} - t_{\text{post}}|}{\tau_d}\right), \quad (20)$$

where $\tau_p$ and $\tau_d$ are the decay constants of LTP and LTD, respectively. Typically, $\tau_p = 40$ ms, $\tau_d = 20$ ms, $C_p = 0.00525$ and $C_d = 0.0105$. Inhibitory weights $G_{ij}$ were modified between zero and an upper bound $G_{\max}$ ($\propto 1/\sqrt{N_{\text{out}}}$).

Evaluation of the degree of independency between signals

ICA was not valid for the auditory signals used for the simulations of BSS. This was because the signals were not independent. In addition to the standard correlations between two analog signals, negentropy (≥0) was used to evaluate the independency of signals. Negentropy measures the deviation of a target distribution from a Gaussian distribution: negentropy vanishes if the target distribution is Gaussian but otherwise takes a positive value; the larger the deviation is, the larger the value of negentropy is. The calculation of negentropy $J(Y)$ for the statistical variable $Y$ requires the true distribution, but it is unknown in the present study. Therefore, we made the following approximation in the evaluation of $J(Y)$ using a certain function $Q$:

$$J(Y) \propto \left[E\big(Q(Y)\big) - E\big(Q(\rho)\big)\right]^2, \quad (21)$$

where $E(x)$ refers to the expectation value of $x$ and $\rho$ obeys a Gaussian distribution. Typically, the logarithm of hyperbolic cosine function is used for $Q$ (*15*):

$$Q(u) = \frac{1}{a} \log\cosh(au), \quad (22)$$

where $1 \leq a \leq 2$. In this study, we set as $a = 1$.

### Supplementary Text

Simulation details

Fig. 2: The numbers of input and output neurons were 1000 and 10, respectively and each input neuron selectively responded to a letter at the firing rate of 10 Hz. In (A) and (B), the individual chunks consisting of four English letters had the same length of 30 ms and appeared in the input sequence with the same occurrence probabilities of 1/3. Letters in each chunk appeared in a fixed order. Both feedforward connections and lateral inhibitory connections were trained.

Fig. 3: The size of each image was 28×28 (= 784) pixels and each bar has a width of 7 pixels. Each pixel took its value on either 1 or 0. Noise was generated by flipping pixels with the probability of 0.1, and the images were blurred by circular masks to prevent artifacts from the edges of the images. Each input neuron received input from an image pixel without overlaps between neurons, and the neurons responding to the pixels of value 1 generated Poisson spike trains with the mean firing rate of 40 Hz. The total time of training was 400 sec. The weights of lateral inhibitory connections were not modifiable.

Fig. 4: The network had 500 input neurons and two output neurons with non-modifiable lateral inhibition. Each input neuron generated a Poisson spike train at an instantaneous firing rate equal to the waveform of the mixture signal assigned randomly to the neuron. During training, the network model was repeatedly exposed to mixture signals 60 times, where each presentation had 500,000 time steps. In sampling spiking activities, we normalized the mixture signals between the minimum (0 Hz) and maximum (10 Hz) rates. Note that, input spike trains varied from trial to trial although their rate profiles were unchanged. The sampling rate of audio files was 44.1 kHz and the unit time step of network simulations was 1 ms. For comparison with ICA, we used the FastICA function of Python library scikit-learn with a log cosh function, and tolerance on update at each iteration was 0.0001 (*8*, *31*). FastICA is efficient and popularly used. Independence of auditory signals was evaluated with the negentropy (negative entropy), which measures the deviations of sampled signals from a Gaussian random process. Correlations between signals were not sensitive enough to discriminate between our model and ICA in performance. We compared our model and ICA in two cases. In the first case, we used the sounds of two music instruments playing different pieces of music. In the second case, we used the sounds of the same two instruments playing their own repertoires of the same piece of music. The summed negentropy of two source signals was 0.001 and 0.0004 for the former and latter cases, respectively, indicating that the two signals were less independent in the latter case than in the former case. Generally, the two source signals were highly correlated. Our model gives output activities that are generally delayed behind the input (Fig. 4D). The delay was adjusted at about 13 ms to obtain the maximal correlations between true sources and output neuron activities. We paired each output activity with the true source that yields the largest correlation. In ICA, the signs of the estimated signals can be opposite to those of the true sources. Therefore, we calculated the correlations between all pairs and all possible combinations of signs (when the number of sources is two, $2! \times 2^2 = 8$ patterns) to adopt the maximal value. In (C), each waveform was reconstructed from the firing rate of each output neuron averaged over 20 trials with an identical set of initial weights after standardization. Namely, we subtracted the temporal average of the (trial-averaged) firing rate from the instantaneous values and divided the resultant differences by the standard deviation. In (D), simulations were performed 400 times for all combinations of 20 different sets of input spike trains and 20 different sets of initial weights.

Fig. S5: In (A), synaptic input to an output neuron during the presentation of a target input pattern was the superposition of a target-specific Poisson spike train of rate $r_{\rm sig} = r/(1 + (S/N)^{-1})$ and a background Poisson spike train of rate $r - r_{\rm sig}$, where $S/N$ refers to the signal-to-noise ratio $r_{\rm sig}/(r - r_{\rm sig})$. Note that $r_{\rm sig} < r$. The temporal patterns of target-specific spike trains were kept unchanged across the repetition of target patterns, whereas background input changed their temporal patterns for each repeat. Outside the target patterns, Poisson spike trains of rate $r$ were given as input. Thus, the spike rate of synaptic input was always $r$, which was fixed at 5 Hz. In (B), we did not model the realistic process of failure in synaptic transmissions, i.e., failure in evoking postsynaptic potentials by presynaptic spikes. Instead, we simulated transmission failure in the following way. We first generated Poisson spike trains with a fixed Poisson rate $r/(1-p_{\mathrm{fail}})$, where $p_{\mathrm{fail}}$ is the failure probability of presynaptic transmissions. Then, assuming that the failure rate is small, we eliminated spikes with the probability $p_{\mathrm{fail}}$. In (C), we introduced trial-by-trial jitters in presynaptic spikes. First, we generated the reference spike trains used for the presentation of a chunk to individual input neurons. Then, in each presentation of the chunk, spike times were sifted by the amounts drawn by a Gaussian distribution with mean zero.

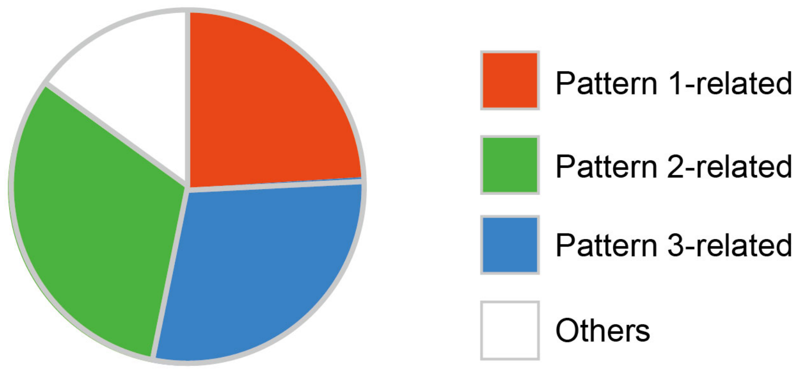


**Fig. S1. Selective learning of repeated spike sequences.** The fraction of trials in which a single dendritic neuron model learned a selective response to one of the three repeated spike patterns is shown. The number of trials was 100. “Others” indicates trials in which the neuron had more than one preferred pattern, i.e., the peak responses to the second preferred pattern was greater than 50% of those to the most preferred pattern.

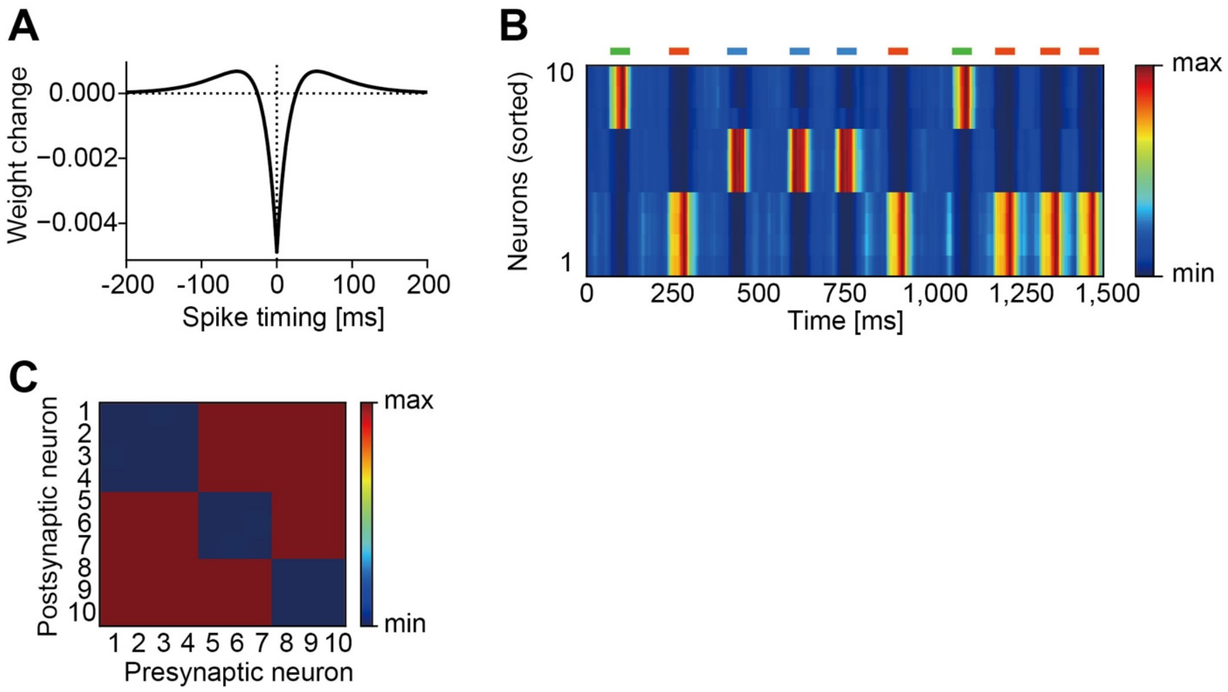


**Fig. S2. Inhibitory STDP.** (A) Window function of the iSTDP implemented at lateral inhibitory connections in the model network. Here, spike timing refers to the time advance of postsynaptic firing from the preceding presynaptic input. The window function of STDP assumed here has been observed at neocortical excitatory synapses on inhibitory neurons (*32*). However, to the best of our knowledge, STDP has not been examined between inhibitory neurons and the present window function is a prediction of our model. (B) Phasic responses of output neurons are shown. Horizontal bars show the intervals in which three chunks (green, red, blue) were presented. The neurons were sorted according to their onset response times to show the emergence of chink-specific cell assemblies. (C) Post-learning synaptic weight matrix of lateral inhibition is shown.

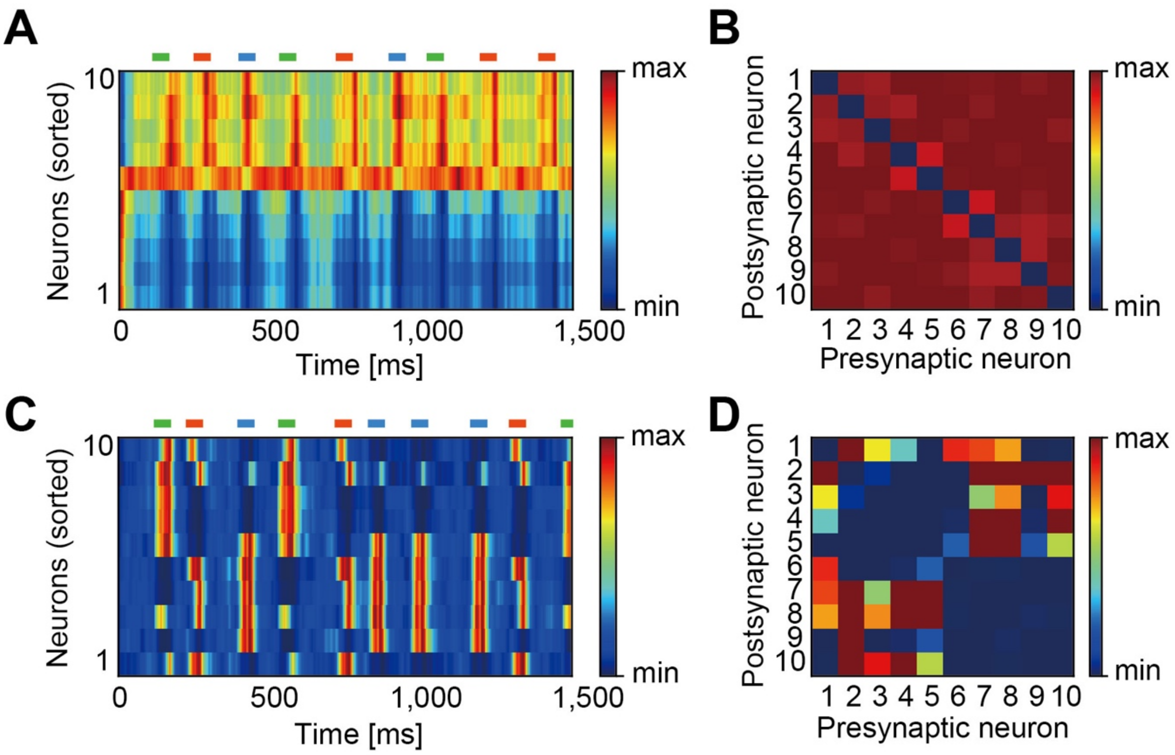


**Fig. S3. Learning with strong or weak lateral inhibition.** The magnitude of lateral inhibition has to be modified in a modest range. Here, simulations were performed with irregular spike trains of 200 input neurons. The other settings were the same as in Fig. S2. (A) Post-learning responses of output neurons are shown for strong lateral inhibition. About half of the neurons responded to all three chunks without stimulus selectivity and the others showed almost no responses. (B) Synaptic weight matrix developed no clustering structures for too strong lateral inhibition. (C) Post-learning responses of output neurons are shown for weak lateral inhibition. The majority of neurons had more than one preferred stimulus. (D) Synaptic weight matrix developed overlapping clustering structures for too weak lateral inhibition.

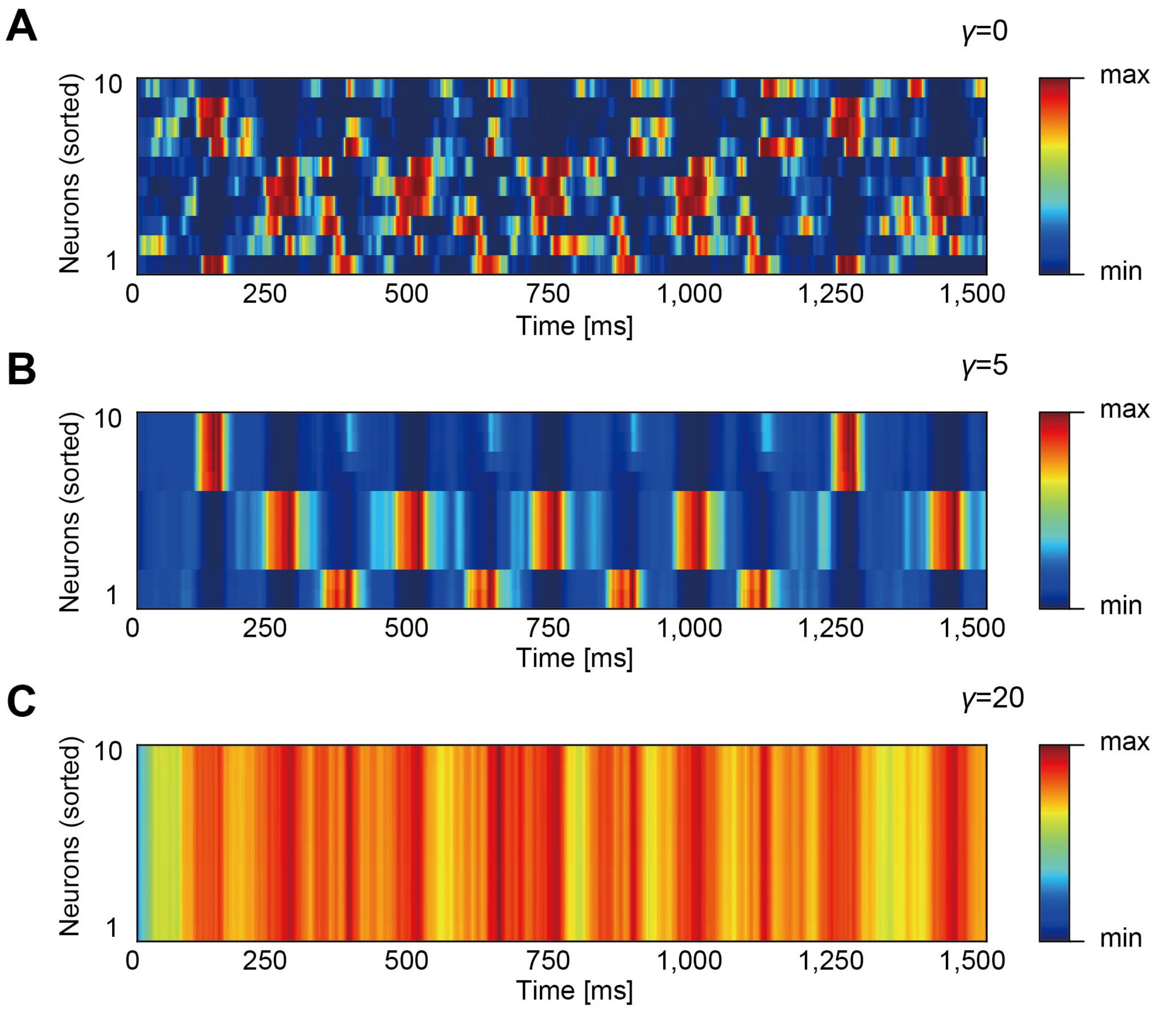


**Fig. S4. The effect of the regularization term on learning.** The strength of the regularization term was changed in Eq. (19) in the task shown in Fig. 2. (**A**) Too weak regularization term impaired the self-organization of selective responses to chunks. (**B**) A moderate range of the regularization term resulted in successful learning. (**C**) Too strong regularization term prohibited the learning of features.

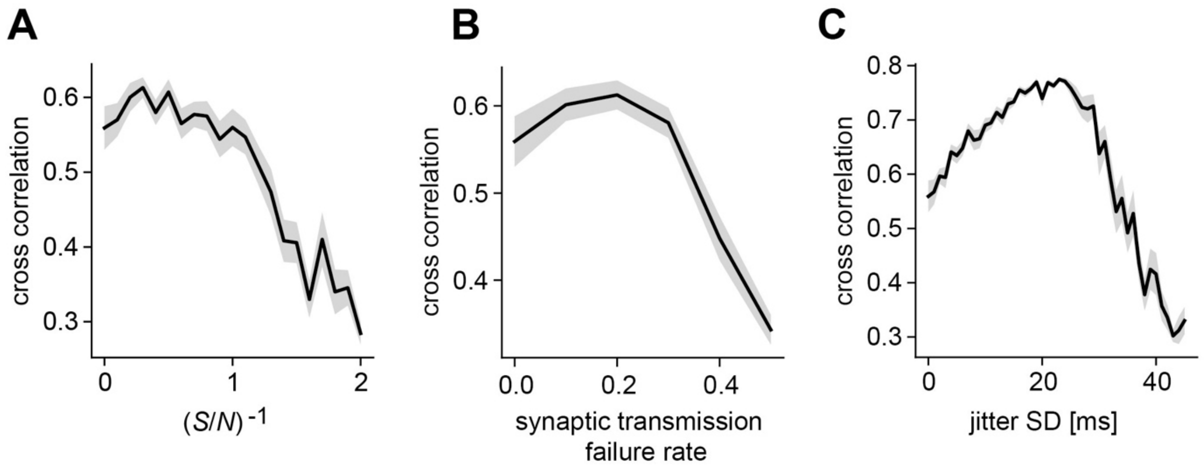


**Fig. S5. Effects of different presynaptic noise.** Performance in learning spatiotemporal patterns of presynaptic spike trains was evaluated for (A) contamination by background presynaptic spikes, (B) failure in synaptic transmissions and (C) jitters in presynaptic spikes in the target activity patterns. The ordinates in (A) and (C) refers to the inverse of the number ratio of background spikes to target-specific spikes and the s.d. of spike timing jitters, respectively. The mean (thick lines) and s.d. (shaded areas) of correlations were calculated between the responses of output neurons and chunk-specific reference responses over 20 trials at each noise level, where a reference response takes unity during the presentation of the corresponding chunk and zero otherwise. We calculated the correlations for all possible pairs of output responses and reference responses, and determined a preferable chunk of each output neuron by the reference response maximally correlated with activity of that neuron. Learning performance was determined as the mean of the maximal correlations over output neurons.

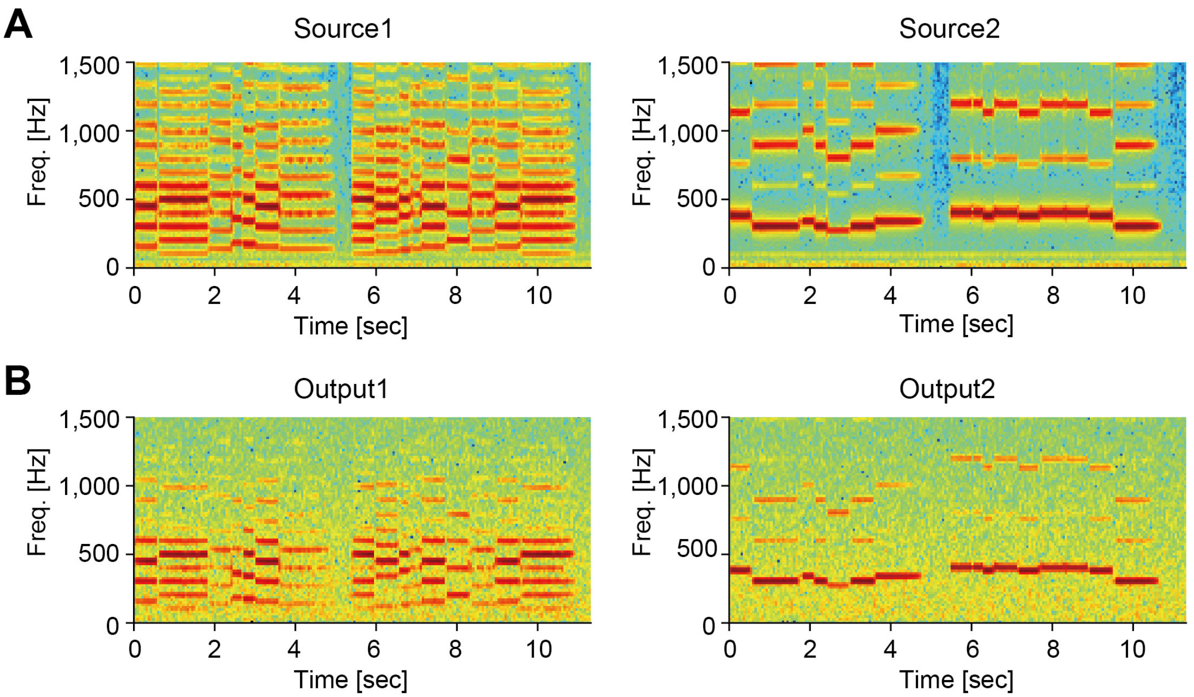


**Fig. S6. Spectrogram of true and estimated signals in BSS.** (A) The spectrograms of the true sources. The two sources were taken from the same piece of music. (B) Example spectrograms of the estimated signals. The network model cut off the high-frequency components above ~ 60 Hz. This was because the membrane dynamics act as a low-pass filter with the cut-off frequency being the inverse of the membrane time constant.

**Audio S1.** An example of mixture sounds of clarinet and bassoon playing their repertoires of an identical piece of music.

**Audio S2.** The original sound of a bassoon**.**

**Audio S3.** The original sound of a clarinet.

**Audio S4.** The sound of bassoon separated by our model.

**Audio S5.** The sound of clarinet separated by our model.

**Audio S6.** One of the signals separated by ICA.
